# Rewilded Highland Cattle in conservation grazing may not groom more when horseflies are present and temperature is higher

**DOI:** 10.1101/2025.01.02.631143

**Authors:** Jinhwi Kim, Pius Korner, Valentin Amrhein, Lilla Lovasz

**Affiliations:** Mammal Team, Research Center for Endangered Species, National Institute of Ecology, Yeongyang, Gyeongbuk Province 36531, Republic of Korea; Research Station Petite Camargue Alsacienne, 68300 Saint-Louis, France; Department of Environmental Sciences, Zoology, University of Basel, CH-4051 Basel, Switzerland; Oikostat GmbH, CH-6218 Ettiswil, Switzerland

**Keywords:** Grooming behavior, Natural behavior, Semi-wild cattle, Highland cattle, Behavioral adaptation

## Abstract

Grooming behavior in domestic cattle serves various functions, including hygiene maintenance, social bonding, and stress alleviation. We examined the grooming patterns of rewilded Highland cattle, to describe their behavioral adaptations and responses to environmental factors in a conservation grazing system. We observed 21 Highland cattle in a French nature reserve from November 2020 to September 2021 using mixed focal and scan sampling methods, recording a total of 1225 grooming bouts. Grooming primarily consisted of self-grooming (83%), followed by tree-grooming (16%), and social-grooming (1%). The average grooming duration per bout was 48.2 seconds per 15-minute interval, indicating a stable and consistent grooming pattern across all observed cattle. We used linear and generalized mixed effect models to assess the effects of environmental factors such as the presence of horseflies, the Temperature-Humidity Index (THI), and habitat type. Our results showed that grooming behavior was influenced by habitat and group, while environmental stressors had only minor effects on grooming duration and frequency. This limited response may be attributed to the relatively low density of horseflies in our study area and the opportunity of adaptive behaviors, such as wallowing, to manage heat and ectoparasites. By examining grooming behavior under near-natural conditions, this study provides a baseline for understanding behavioral patterns and adaptations in rewilded cattle, while also serving as a potential reference for identifying behavioral changes in domestic cattle and informing future management practices.

## Introduction

Grooming in domestic cattle serves multiple functions, including removing foreign objects (Hart, 1990), maintaining hygiene (Spruijt et al., 1992), establishing social relationships (De Freslon et al., 2020; Reinhardt et al., 1986; Sato et al., 1991), and developing maternal bonds (Kohari et al., 2009; Newby et al., 2013). It is also a comfort behavior that helps animals cope with stress (Park et al., 2020; Spruijt et al., 1992), such as social isolation (Mandel et al., 2019), calf separation (Newby et al., 2013), or novel neighbors (Herskin et al., 2004), particularly in intensive farming systems. In contrast, in extensive or semi-wild environments, where stressors related to husbandry are largely absent, grooming may serve alternative functions, such as managing ectoparasites or maintaining self-care, reflecting behavioral adaptations to natural surroundings.

Cattle in domestic setups have been shown to have a strong desire for grooming. For instance, a push-gate experiment (McConnachie et al., 2018) showed that the motivation to use a grooming brush was as strong as the motivation to access fresh food; and in another experiment, cattle used a brush regularly despite skin issues (Moncada et al., 2020). However, when cattle were restricted from grooming due to the lack of appropriate grooming substrates and small living areas, they tended to engage in intensive grooming with short bouts, often scratching themselves against barn facilities (Anselme, 2008). Consequently, some studies have suggested using grooming behavior as a potential on-farm metric of domestic animal welfare (Napolitano et al., 2009; Winckler et al., 2009) and as a health monitoring index (Mandel et al., 2019; Toaff-Rosenstein et al., 2017).

However, there are practical challenges in employing grooming as a reliable indicator of cattle welfare. First, despite increased interest in pasture-based, animal-friendly systems (Moscovici Joubran et al., 2021; Stampa et al., 2020), animal-based measures for assessing the welfare of ruminants in extensive and semi-wild systems remain underexplored (Spigarelli et al., 2020): there are only a few studies that examined the grooming pattern of cattle in extensive housing and semi-wild conditions (Hall, 1989; Krohn, 1994; Reinhardt et al., 1986). Second, previous grooming studies have focused on brush use in indoor-housed cattle, but the comparability of the results of these studies are limited due to the distinct methodologies used (i.e., varying time-sampling methods, durations, and observation periods) across studies (Horvath et al., 2020). Lastly, because indoor-housed cattle are usually dehorned and their hooves are trimmed by owners, grooming activities associated with these parts are often excluded from quantification and remain unknown. Investigating grooming behavior in natural or near-natural conditions could therefore lead to a more reliable baseline measure which can then be used as a reference for reliably using grooming behavior as a welfare index in domestic conditions.

Extensive and semi-wild conditions expose animals to a variety of environmental stimuli (Mellor, 2015), but also pose multiple challenges (e.g., parasites, variable climate, and predation). Therefore, animals in such environments may adapt their behavioral patterns, leading to changes in their time budget, and the frequency and duration of behaviors in their repertoires (Hart, 1990). For example, Krohn (1994) reported that cattle altered their grooming patterns (i.e., frequency, duration, and targeted body parts) when they were allowed to access extensive grazing areas. This suggests that cattle may adjust their grooming patterns in response to novel stimuli and other environmental factors associated with different living conditions (Hart, 1990; Kohari et al., 2007). Grooming may thus be a response to natural stressors such as insect harassment or high temperatures, but it may also indicate social behavioral responses or reactions to certain environmental or habitat variables; however, to our knowledge, no study yet addressed this topic in natural or near-natural conditions.

Our study therefore aimed to quantify and characterize the grooming patterns of rewilded (Lovász et al., 2024) Highland cattle – a rustic breed, well adapted to natural conditions – in a French nature reserve. We also investigated the influence of environmental stressors, including the presence of horseflies, temperature fluctuations, and the effect of season and different habitats, on grooming behavior. Our expectations were the followings:

1. During the summer months, the presence of horseflies will increase grooming frequency but decrease grooming duration, as cattle may engage in rapid, single-stroke grooming to respond to horsefly attacks.
2. High ambient temperatures will reduce both the frequency and duration of grooming behavior, as the energy expenditure associated with grooming may become costly under hotter conditions.

## Materials and methods

### Study site

The study was conducted from November 2020 to September 2021 in in the national nature reserve Petite Camargue Alsacienne in Saint-Louis, France. The climate in this area is typically moderate, with warm summers and cold winters. The mean ambient air temperature was 11.15°C and temperatures ranged from -13 to 34.6°C for the study period, based on data from the local weather station (“Weather Forecast for Rosenau, France,” 2022). The approximately 47-hectare cattle enclosure (**Fig. 1**) comprises a mosaic of wet and dry environments, including swamps, reedbeds, marshes, shrub- and forest-covered areas, and meadows.

**Figure 1.**
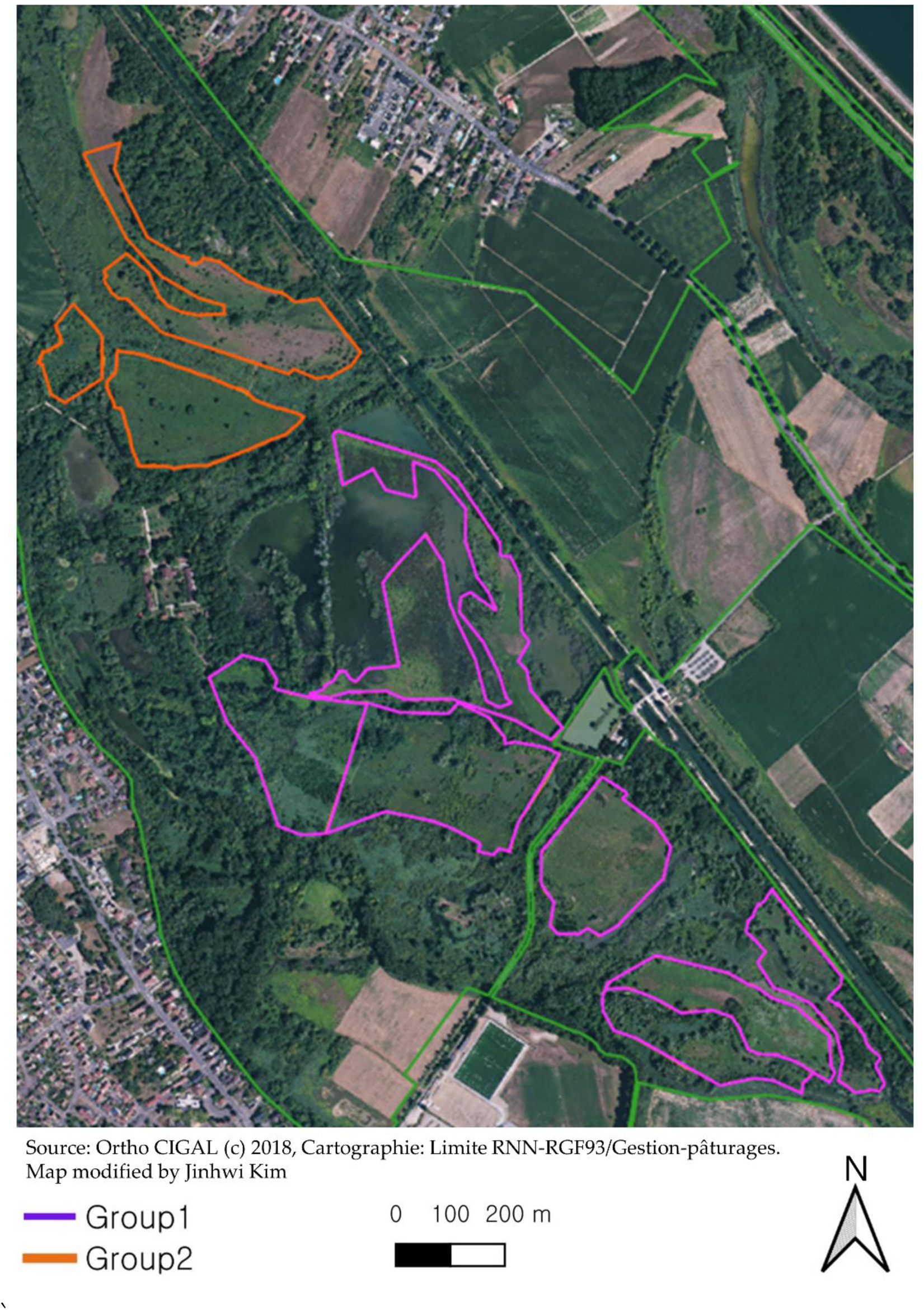
Map of the study site. The satellite image shows the locations of the two distinct enclosures, marked in orange and purple, where the two groups of Highland cattle remain year-round.

### Animals and data collection

Highland cattle, characterized by long horns, double hair coats, and relatively small body size, are particularly adapted to thriving in harsh environments and cause minimal damage to swards and soft soils (Tolhurst, 2001). Due to these characteristics, Highland cattle have been introduced into various European nature reserve areas with diverse conservation grazing conditions (Lamoot et al., 2005). Our study animals were two herds of Highland cattle in the Petite Camargue Alsacienne that graze in the nature reserve year-round with minimal human intervention.

At the beginning of the observation period in November 2020, 25 Highland cattle were selected as study subjects, varying in sex and age. During winter, however, three cattle died of parasitic infections caused by a common liver fluke (E. Linder, personal communication), and one cow was excluded from the study following the management plan in June 2021, resulting in 21 animals in total. Group 1 consisted of 11 individuals (8 steers and 3 cows), with an average age of 11.1 ± 3.6 years (range: 7–16 years), whereas group 2 consisted of 10 individuals (2 steers and 8 cows), with an average age of 6.3 ± 4.2 years (rage: 2–16 years). The two groups were rotated between the several sub-enclosures based on the management plan and grass conditions. During the winter feeding period, the herds stayed in a sub-enclosure equipped with a hay rack, where supplementary fodder (dry hay) was provided upon necessity (e.g., depleted resources or long-term snow cover).

### Behavioral observation

We conducted observations 2-3 days per week. Field observations were conducted under favorable weather conditions to minimize the influence of precipitation or heavy wind on cattle behavior. We used focal and scan sampling methods (Altmann, 1974; Crockett & Ha, 2010) to record individual grooming patterns, time budgets, and habitat use of all group members. Each observation day consisted of three 65-mintue observation sessions conducted during daytime: morning (07:30-08:35 h), noon (11:30-12:35 h), and afternoon (16:00-17:05 h). The observation times were set according to the diurnal rhythm of pasture-raised cattle (Kilgour, 2012), which peaks around sunrise and sunset for grazing.

Each observation session was subdivided into four 5-min scan sampling periods and three 15-min focal sampling periods. The 5-min scan sampling periods were designed to accommodate both the brief time needed to record behaviors and any additional time required to locate all individuals in the herds. Observations were conducted sequentially for the two cattle herds, and focal animals were preselected by shuffling individual animal numbers before each observation session, thus systematically randomizing the sampled individuals while avoiding sampling the same individual twice in one session. During scan sampling, the observer (J.K) recorded the habitat where the majority of group members were located and noted the behaviors of all group members based on an ethogram developed for scan sampling (**Table 1**). During focal sampling, the following variables were recorded according to an ethogram for focal sampling (**Table 1**): grooming types, targeted body parts, the habitat where the cattle displayed grooming (**Table 2**), and the grooming duration of the focal animal. The ethograms and variables were tailored based on frameworks suggested by Kilgour (2012) and Krohn (1994), to quantify and characterize grooming patterns effectively, capturing group-level behaviors during scan sampling and individual-level grooming details during focal sampling. Grooming behavior in cattle, as observed in our study, typically follows pre-grooming behaviors such as pausing, exhibiting vigilance, turning the neck toward the grooming area, and salivating. A grooming bout was defined as a continuous grooming sequence directed at the same body part. If there was no continuous grooming within 10 s, or if the animal started to groom a different body part, it was considered a separate grooming bout.

**Table 1.**
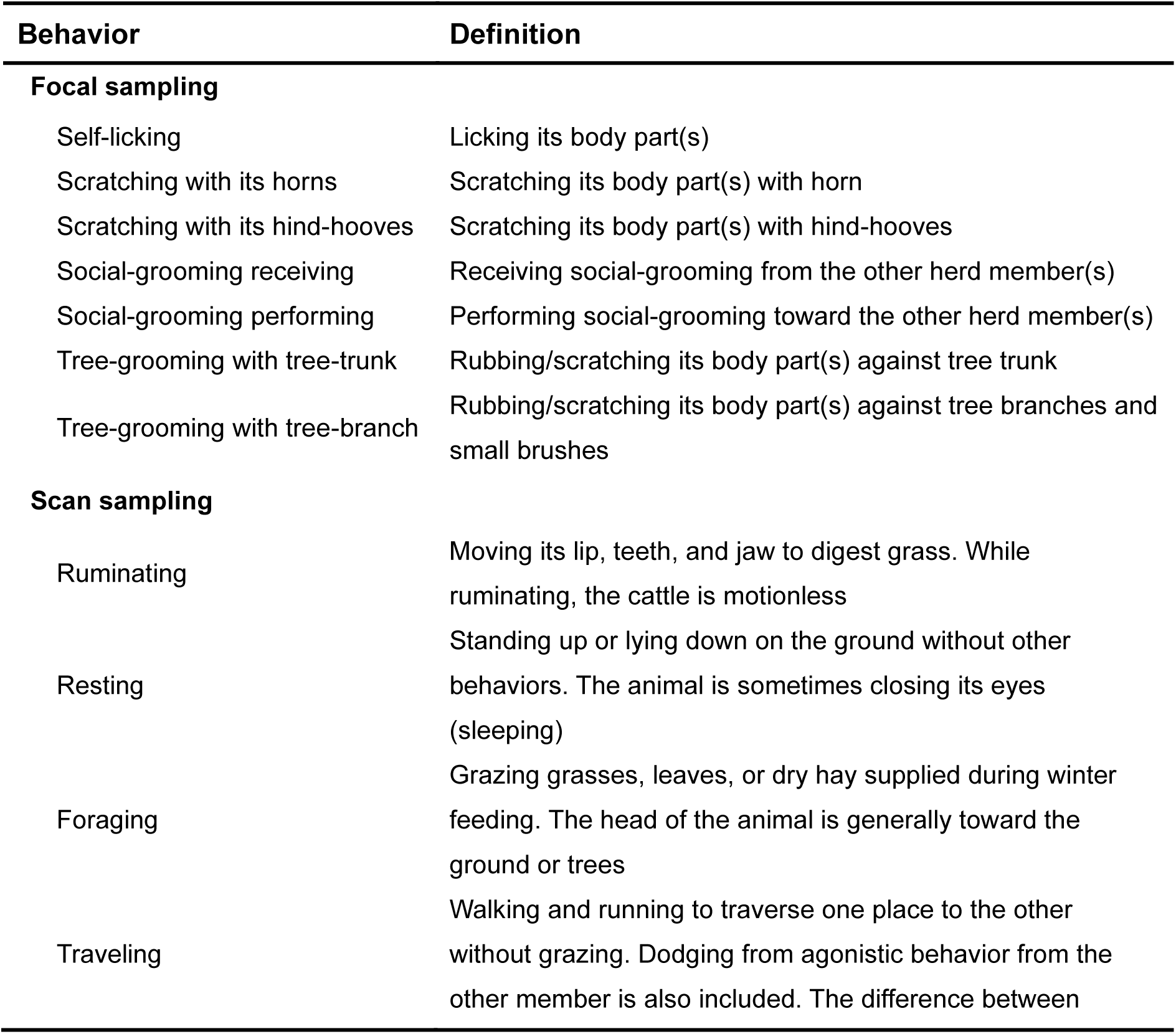
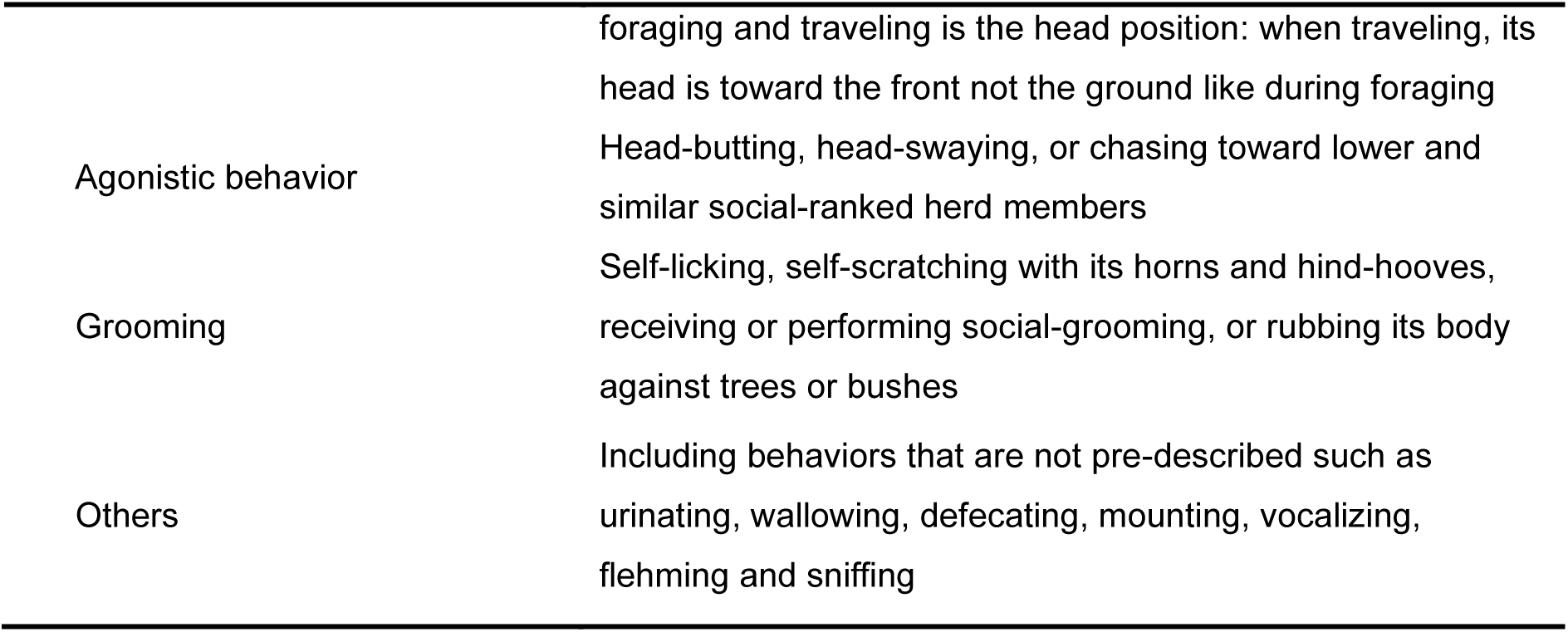
Ethogram of the behavior of semi-wild cattle for focal and scan sampling. This table outlines the behavioral categories and definitions used during mixed focal and scan sampling. The ethogram was tailored to effectively capture both individual-level and group-level behaviors of Highland cattle in semi-wild conditions, drawing on established frameworks by Kilgour (2012) and Krohn (1994). Adaptations were made to ensure precise quantification and characterization of grooming patterns and other key behaviors.

**Table 2.** Description of variables. Variables used to analyze the context and patterns of grooming behavior in Highland cattle. Body part variables refer to specific anatomical regions targeted during grooming, such as the front, back, and inner body. Habitat variables describe the environments where behaviors occurred, including meadow, forest, swamp, and feeder areas.

| Variables | Definition |
| --- | --- |
| **Body parts** |  |
| Head | Including forehead, face, nose, and muzzle |
| Neck | Including throat, neck, dewlap, and brisket |
| Front | Including front legs, shoulders, elbows, foreflank, and front part of body |
| Back | Including hind legs, back, loin, rump, and tail |
| Inner part | Including udder, teats, belly, sheath, and testicles |
| **Habitat** |  |
| Meadow | A small or medium sized grassy area with trees and bushes |
| Forest | An area of tree-covered land where the canopy cover is more than 20% |
| Marshes | Seasonal freshwater marshes/pools on inorganic soils<br>Including seasonally flooded meadows, sedge marshes, shrub-swamp, and reedbeds |
| Hay rack (feeder) | A place for storing hay and feeding herd during winter feeding |

### Statistical analysis

To investigate the impact of horseflies, we selected grooming data from the summer months (125 bouts from 42 focal sampling sessions in June, July, August, and September 2021), when horseflies were active at the study site. To estimate environmental heat exposure, we calculated the Temperature-Humidity index (THI) using the “weathermetrics” R package, as suggested by Anderson et al., (2013). THI is a combined measure of ambient temperature and humidity, used to assess the overall heat stress experienced by animals in a given environment. In this study, we used THI to evaluate the potential impact of heat stress on the grooming behavior of Highland cattle in semi-wild conditions. Because cattle sometimes moved between habitats during sampling, we assigned the first recorded habitat as the primary habitat for analysis.

We used two models to estimate effects of the presence of horseflies, THI, day of the year, habitat, group, and age of animals on the average grooming duration and frequency, respectively. The log-transformed grooming duration was modelled using a Gaussian data distribution, and the frequency of grooming during 15 minutes of observation using a Poisson data distribution and log-link function. We z-transformed the predictor variables. Since the day of the year and age of animals showed a non-linear relationship with frequency, we used a linear and quadratic polynomial to fit the curve. The individual animal and the observation session was used as a random factor.

We fitted the model using a Bayesian approach via the rstanarm package (Stan development team, 2022) and the brms package (Bürkner, 2017) using default priors and settings. Model checking was performed using QQ plots of the residuals, checking for overdispersion in the Poisson model, plotting residual vs. predictors, and using posterior predictive model checking. Medians of the marginal posterior distributions of the model parameters were used as estimates, and the 2.5% and 97.5% quantiles were used as the lower and upper limits of the 95% compatibility intervals (Amrhein and Greenland, 2022). All analyses were performed using R Statistical Software (v4.2.2; R Core Team 2022)

## Results

### Characterization of cattle grooming

We conducted 55 field observation sessions and observed 1225 grooming bouts. The overall grooming duration per single bout ranged from 0 to 592 seconds per 15-minute interval, with an average grooming duration of 48.2 seconds per 15-minute focal sampling. The mean grooming durations for different grooming types (**Table 1**) were as follows (**Fig. 2**): self-grooming (41.5 s / 15 min), tree-grooming (65.8 s / 15 min), social-grooming (106.4 s / 15 min). The mean values were relatively higher than the median durations (self: 21.4; tree: 28.5; social: 90.4 s / 15 min), due to the presence of extreme outliers.

**Figure 2.**
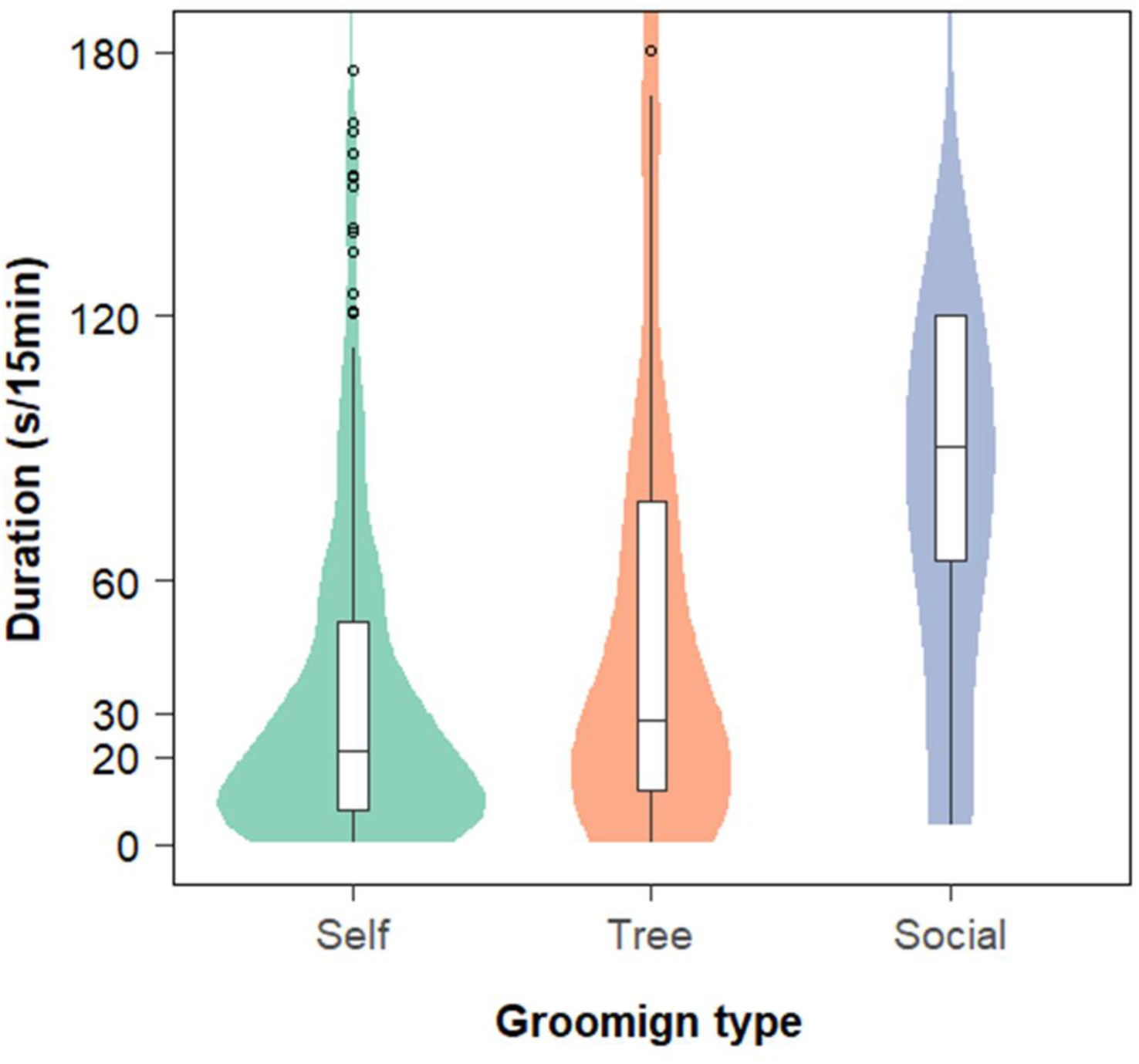
Grooming duration per focal sampling across different grooming types. Violin plots and box plots depict the average grooming duration per 15-min focal sampling period (Self: n = 1017; Tree: n = 194, Social: n = 14). Extreme outliers below 180 seconds are shown as blank circles, while n = 7 outliers exceeding the y-axis range are not displayed in the figure.

Self-grooming accounted for 83% of the total grooming bouts, followed by tree-grooming (16%) and social-grooming (1%) (**Fig.3**). Grooming frequency was consistently observed throughout the day, with a similar distribution across the morning (33%), noon (32%), and afternoon (36%) periods. Among the various grooming sub-behaviors, self-licking (48%) was the most frequently displayed, followed by horn scratching (28%), scratching against tree branches (12%), hind-hoof scratching (7%), scratching against tree trunks (4%), and social grooming (1%). Out of the five distinguished body areas, the front part was groomed the most frequently, followed by the back, head, neck, and inner body parts.

**Figure 3.**
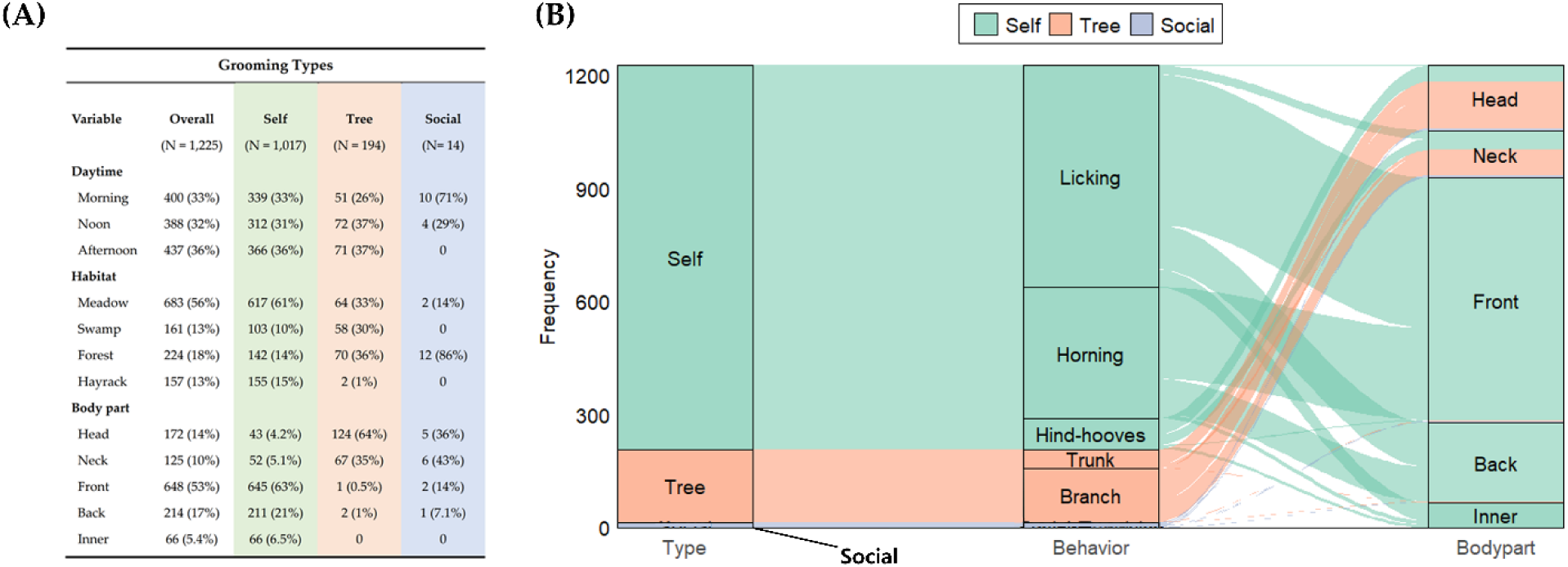
Relationships between variables and grooming types. (A) illustrates the grooming patterns of Highland cattle characterized by grooming types, daytime, habitats, and body parts. The alluvial plot (B) shows relationships between grooming types, grooming sub-behaviors, and body parts (n = 1,225 grooming bouts). The height of strata is proportional to the grooming frequency. Social grooming was rare (n = 14) and is hardly visible in the plot.

We collected 5734 scan data points from the two groups of Highland cattle based on the ethogram for scan sampling. Three survival-related behaviors (foraging, resting, and ruminating) accounted for 84.3% of the total behavioral repertoires (**Fig. 4(A)**). Both herds consistently engaged in grooming throughout the year, allocating approximately 10% of their diurnal time budget to this activity (**Fig. 4(B)**). Agonistic interactions among group members were rare (0.2%) and were primarily observed during the winter feeding time.

**Figure 4.**
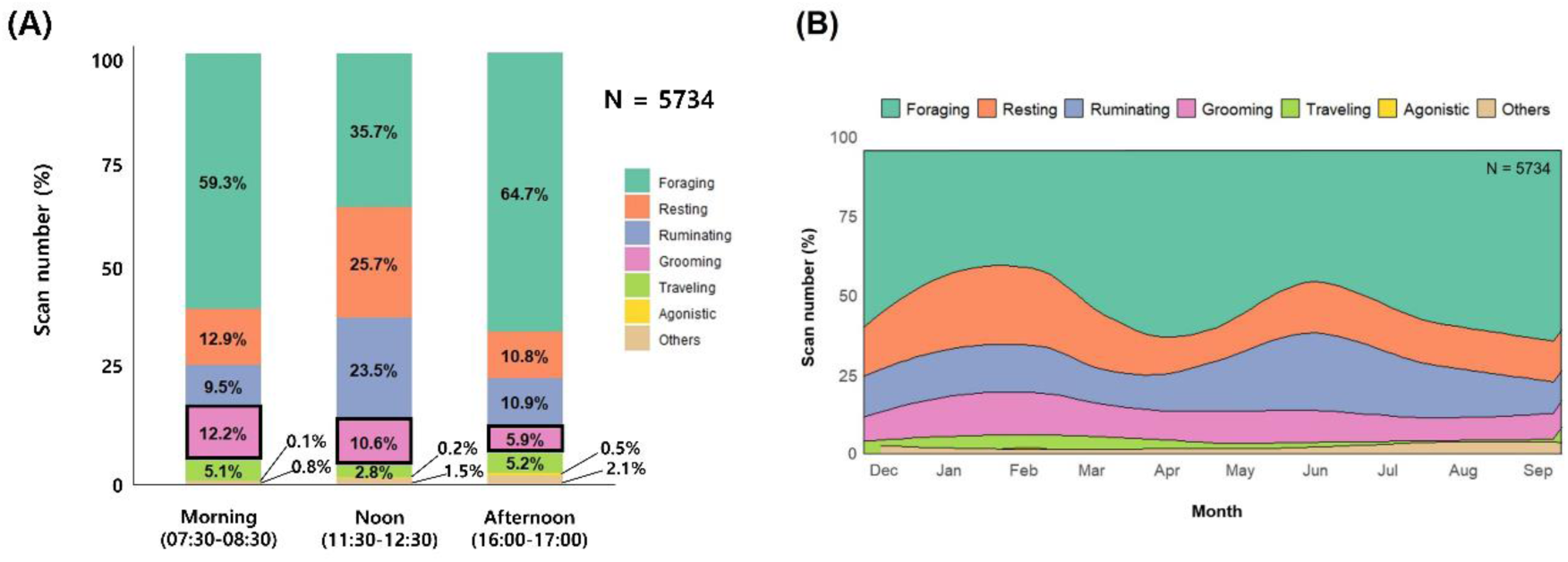
Diurnal behavior patterns by different times of day and months. (A) shows the percentage of scan data (morning: n = 1929; noon: n = 1866; afternoon: n = 1939) recorded during different times of the day. ‘Agonistic’ and ‘Other’ behaviors were very rare (n= 14, n = 84) and are hardly visible. Grooming is highlighted with solid lines. (B) shows the behavior patterns of two groups of cattle from December 2020 to September 2021.

### Effect of environmental factors

Throughout the study period, horseflies were only present from June to September 2021. During a 15-minute session, a maximum of three flies were observed around an individual. Although most tabanid fly species are usually present for only about one month of the year, their succession throughout the warm months leads to cattle being exposed to attacks by horseflies across the entire warm season (Foil and Hogsette, 1994). During this same period, the average temperature was 21.4°C (70.3°F), with an average humidity of 65%. The corresponding THI was 70.3.

Overall, the effects of external stressors, i.e., the presence of horseflies and temperature and humidity as measured by the Temperature-Humidity Index (THI), showed no strong effects on grooming duration and frequency in comparison with other variables (**Fig. 5**). Grooming duration slightly decreased when horseflies were present, while grooming frequency was similar, but uncertainty was high (CIs were wide; **Fig. 5 and 6**).

**Figure 5.**
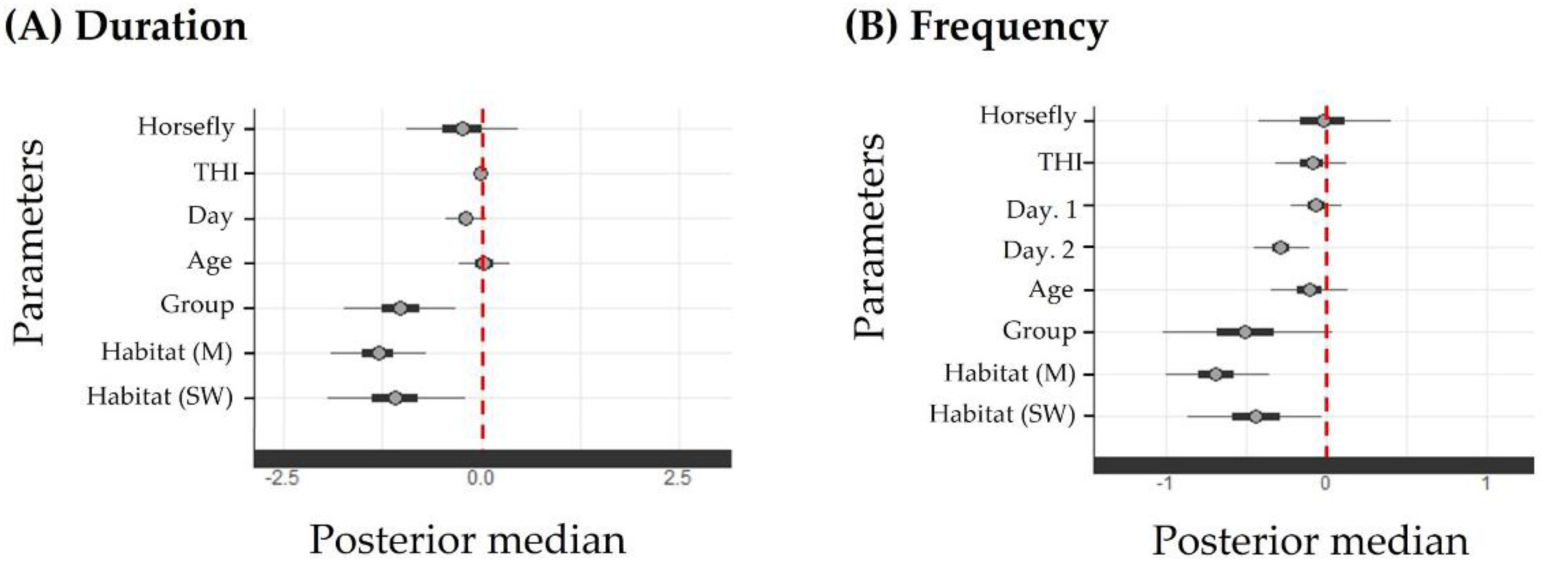
Estimated coefficients for grooming duration (A) and grooming frequency (B) in response to various external stressors as well as age, group and habitat. The x-axis shows values of the marginal posterior distributions. The red dashed line at zero indicates no effect. Thicker bars are 50% Bayesian compatibility intervals (CI), capturing values that are most compatible with the data and the model, while thinner bars are 95% Bayesian compatibility intervals. “THI” is Temperature-Humidity Index. In panel (A), “Day” represents the day of the year, while in panel (B), “Day. 1” and “Day. 2” denote the linear and quadratic terms, respectively, from orthogonal polynomials used to model non-linear effects of the time of year. “Group” is one of the two studied herds of cattle against the other, “Habitat M” is meadows and “Habitat SW” is swamp habitat.

**Figure 6.**
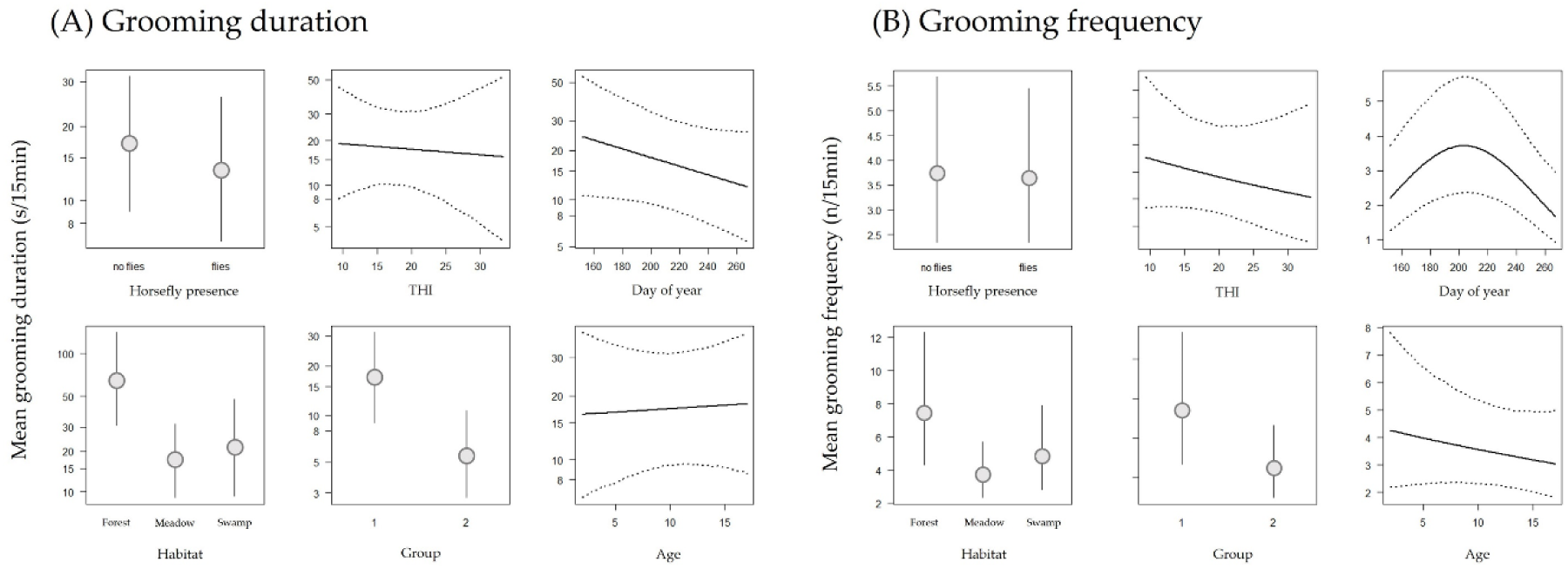
Effect plots of (A) grooming duration (s / 15min) and (B) frequency (n / 15min) in relation to horsefly presence, Temperature Humidity Index (THI), day of year, habitat type, group number and age of animal. Given are point estimates (circles) and regression lines with 95% Bayesian compatibility intervals of the marginal posterior distributions.

Uncertainty was lower (CIs were shorter) for THI, day of the year, and age: we found virtually no effect of animal age on grooming duration (**Fig. 5A)**, even though grooming frequency showed a slight decline in older animals (**Fig. 6B**). There was only a slight negative effect of day of year on grooming duration (**Fig. 5A**), meaning that grooming duration slightly decreased during the summer months (**Fig. 6A**). There was a clear peak of grooming frequency in the middle of the summer in July (**Fig. 5B and 6B)**.

Grooming duration and frequency were clearly lower in Group 2 compared to Group 1 and for meadow and swamp habitats compared to forest (**Fig. 5A and 6A**). As already mentioned, Group 1 included older animals and a higher number of steers, whereas Group 2 included younger animals and a higher proportion of cows; further, although both groups had similar access to habitat types, Group 1 had a larger overall range (as described in Fig. 1).

## Discussion

The present study explored the grooming patterns of rewilded Highland cattle introduced as actors of conservation grazing in a nature reserve. We observed that the cattle consistently engaged in grooming throughout the year and showed specific patterns in grooming types and targeted body area. Unlike farm cattle (Phillips and Rind, 2002; Sato et al., 1991) and feral cattle (Hodgson et al., 2024), which frequently engage in social grooming, our Highland cattle in near-natural conditions rarely groomed each other. This observation aligns with findings by Hall (1989) in Chillingham cattle under semi-wild conditions. The relatively small body size and large horns of both breeds enable self-grooming of nearly all body parts, thereby probably reducing the need for social contact. Furthermore, abundant grooming substrates in natural environments (Meneses et al., 2021) and a rigid social hierarchy in semi-wild settings (Reinhardt et al., 1986) might reduce the necessity for social grooming. This behavioral adjustment may partly explain more varied time allocation patterns in survival-related activities. Survival-related behaviors, such as foraging, resting, and ruminating, accounted for 84.3% of the total behavioral repertoire in this study, which is 5-10% lower than previously reported time budget for cattle in environments with little human interference (Kilgour, 2012). This lower proportion of survival-related activities may suggest that the time saved may be reallocated to behaviors like grooming, often described as a luxury activity (Mandel et al., 2013) in semi-wild settings.

Despite the expectation that environmental stressors such as ectoparasites and temperature would affect grooming behavior (Villalba et al., 2016), our results showed minor effects of these variables on grooming frequency and duration. For example, horseflies known for their painful bites and potential to cause production losses in intensive setups by reducing grazing time (Baldacchino et al., 2014; Foil and Hogsette, 1994; Perich et al., 1986) were expected to increase grooming frequency and decrease grooming duration, because cattle usually perform quick, single-stroke grooming to alleviate bites (Mooring et al., 2004; Mullens, 2019). However, we did not observe noticeable changes in grooming behavior in response to horseflies, likely due to the relatively low density of horseflies in the study area. Foil and Hogsette (1994) reported that horsefly densities of 66-90 flies per animal per day resulted in weight loss of 0.08-0.10 kg per animal per day, suggesting that higher densities are required to cause severe disruption. Similarly, Roberts and Pund (1974) found that a fly population of 2.7 to 8.4 flies per 2-hour period per animal can disrupt the weight gain of domestic cattle. Alternatively, grooming may not be the primary anti-horsefly behavior for Highland cattle, for example if sufficiently deep water ponds are available for wallowing (i.e. standing in water), to deter ectoparasites, or if forested areas are available for shelter (e.g., horseflies are less active in shaded areas (Horváth, 2024)). Horseflies tend to favor landing on the legs of cattle (Mullens, 2019), which are less protected by long hair compared to other body parts, making them more vulnerable to bites. This vulnerability might drive cattle to engage in wallowing or staying in shade, but as our sampling method did not specifically account for these behaviors, further research is needed to quantify their role in ectoparasite defense in semi-wild conditions.

We also anticipated that high temperatures and humidity, as indicated by the Temperature-Humidity Index (THI), during summer would reduce grooming duration, as increased thermoregulatory costs could shift cattle’s behavioral priorities to feeding and resting (Bernabucci et al., 2014; Gaughan et al., 2009). Nonetheless, we found no marked effect of THI on grooming. One possible explanation is that Highland cattle may rely on behavioral adaptations such as wallowing in water and staying in shaded areas – when available – to help stabilize body temperature under heat stress. This behavior, previously noted as a strategy to deter horseflies, might also serve a thermoregulatory function, allowing cattle to maintain consistent grooming patterns without further adjustments in response to thermal stress.

Similarly, the observed grooming behavior in Highland cattle was not strongly age-dependent. Grooming durations were fairly consistent across all age groups, while grooming frequencies showed a slight tendency to be higher in younger animals. However, the compatibility intervals overlapped substantially across age groups, providing weak support for this trend. This uniformity in grooming duration may reflect the breed’s ability to self-groom effectively across all life stages, likely due to their distinctive traits such as small body size and large horns, which enable thorough self-maintenance. The slight increase in grooming frequency among younger animals may align with the body-size principle, which suggests that individuals with a higher surface area-to-mass ratio are more susceptible to ectoparasites (Hart, 1990; Mooring et al., 2000). However, our study animals were, on average, 9 years old, with only 6 individuals younger than 5 years. Highland cattle are known to grow gradually until about 5 years of age, after which they typically maintain a stable body size (Pauler et al., 2019). This skewed age distribution, with most individuals being older, may have reduced variation in body size across the group, potentially limiting the expression of the body-size principle in this study. Future studies with a broader age distribution, or specifically including calves and younger cattle in semi-wild conditions, would be valuable in clarifying the role of age and body size in shaping grooming behavior.

We observed strong habitat and group effects on grooming behavior. In particular, grooming duration and frequency were higher in forested areas, likely due to the abundance of trees serving as convenient substrates for tree-grooming. While tree-grooming frequency was similar between forest and meadow, the higher prevalence of self-grooming might have diluted the overall impact of tree-grooming on grooming behavior in the meadow. Kohari et al. (2007) observed a similar phenomenon in domestic cattle, where providing trees experimentally increased grooming activity. Dickson et al. (2024) further demonstrated that the loss of grooming substrates led to reduced social grooming and grooming directed at other objects, reflecting reduced overall welfare in grazing cattle. Our study extends these findings to semi-wild conditions, emphasizing how natural habitat features shape grooming patterns.

Group-level differences in grooming duration and frequency were also observed, with Group 1 grooming more frequently and for longer durations than Group 2. Although the groups differed in composition, we did not find a strong effect of sex on grooming behavior, and this variable was subsequently removed from the statistical analysis. Both groups had access to similar habitat types, suggesting that habitat differences may not have strongly influenced grooming behavior. Interestingly, Group 1 displayed longer and more frequent grooming near swamp areas, which are typically associated with shorter and less frequent grooming. This unexpected result underscores the potential influence of group-specific dynamics or unmeasured factors on grooming patterns.

Overall, our findings suggest that the grooming behavior in rewilded Highland cattle remains relatively stable and consistent component of their behavioral repertoire in semi-wild conditions, at least at moderate temperatures and numbers of horseflies. Their ability to perform self-grooming across various body parts and the availability of natural substrates may contribute to maintaining a clear social hierarchy, thus minimalizing social grooming and agonistic behaviors. Interestingly, this consistency in grooming behavior is in contrast with the more variable patterns often observed in domestic cattle, which may have welfare implications (Dickson et al., 2024; Park et al., 2020). For example, under certain circumstances in intensive farming systems, grooming may shift from its natural functions – such as hygiene and ectoparasite management – to coping mechanisms in response to chronic stressors (Krohn, 1994; Meneses et al., 2021). However, it is important to acknowledge that farming systems vary, and well-managed systems that allow for more natural behaviors can enable grooming to fulfill its intended roles without leading to maladaptive responses (Meneses et al., 2021; Park et al., 2020; Tuomisto et al., 2008). By describing the grooming behavior of cattle in near-natural conditions, this study provides a valuable baseline for assessing behavioral deviations in domestic settings, contributing to future discussions on welfare standards for rewilded and domesticated cattle.

## Declarations

## Acknowledgements

We thank the team of the Réserve Naturelle Petite Camargue Alsacienne for making it possible to conduct our research in the nature reserve.

## Conflicts of interest/Competing interets

The authors declare that they have no known competing financial interests or personal relationships that could have appeared to influence the work reported in this paper.

## Ethics approval

Our study was purely observational and did not involve interactions with or experiments on animals; it therefore did not require ethics approval.

## Availability of data and code

The data and code are available at OSF: Kim, J., Korner, P., Amrhein, V., & Lovász, L. 2024. “Rewilded Highland Cattle in conservation grazing may not groom more when horseflies are present and temperature is higher.” OSF. December 24, 2024 doi: 10.17605/OSF.IO/ZNRK2

## Authors’ contributions

JK: Conceptualization, Investigation, Data curation, Formal analysis, Original draft, Writing, Review-editing. PK: Formal analysis, Writing, Review-editing; VA: Supervision, Conceptualization, Writing, Review-editing; LL: Supervision, Conceptualization, Investigation, Formal analysis, Writing, Review-editing.

